# SynaptoPAC, an Optogenetic Tool for Induction of Presynaptic Plasticity

**DOI:** 10.1101/2020.03.27.011635

**Authors:** Silvia Oldani, Laura Moreno-Velasquez, Alexander Stumpf, Christian Rosenmund, Dietmar Schmitz, Benjamin R. Rost

## Abstract

Optogenetic manipulations have transformed neuroscience in recent years. While sophisticated tools now exist for controlling the firing patterns of neurons, it remains challenging to optogenetically define the plasticity state of individual synapses. A variety of synapses in the mammalian brain express presynaptic long-term potentiation (LTP) upon elevation of presynaptic cyclic adenosine monophosphate (cAMP), but the molecular expression mechanisms as well as the impact of presynaptic LTP on network activity and behavior are not fully understood. In order to establish optogenetic control of presynaptic cAMP levels and thereby presynaptic potentiation, we developed synaptoPAC, a presynaptically targeted version of the photoactivated adenylyl cyclase bPAC. In cultures of hippocampal granule cells, activation of synaptoPAC with blue light increases action potential-evoked transmission, an effect not seen in hippocampal cultures of non-granule cells. In acute brain slices, synaptoPAC activation immediately triggers a strong presynaptic potentiation at mossy fiber terminals in CA3, but not at Schaffer collateral synapse in CA1. Following light-triggered potentiation, mossy fiber transmission decreases within 20 minutes, but remains enhanced still after 30 min. Optogenetic potentiation alters the short-term plasticity dynamics of release, reminiscent of presynaptic LTP. SynaptoPAC is the first optogenetic tool that allows acute light-controlled potentiation of transmitter release at specific synapses of the brain, and will enable to investigate the role of presynaptic potentiation in network function and the animal’s behavior in an unprecedented manner.

**Significance Statement:** SynaptoPAC is a novel optogenetic tool that allows increasing synaptic transmission by light-controlled induction of presynaptic plasticity.

## Introduction

Long-term plasticity of synaptic transmission is considered as the cellular basis of learning and memory (Takeuchi et al., 2014). Specific patterns of activity can either strengthen or weaken synapses on timescales ranging from minutes to days, thereby shaping the functional connectivity of a neuronal ensemble. While postsynaptic plasticity involves changes in the number of receptors (Herring and Nicoll, 2016), presynaptic plasticity alters the transmitter release probability or the number of release sites (Monday et al., 2018). Presynaptic long-term potentiation (LTP) is initiated by an increase in cyclic adenosine monophosphate (cAMP) produced by calcium-stimulated adenylyl-cyclases in the synaptic terminal (Ferguson and Storm, 2004). This was first described for hippocampal mossy fiber (MF) synapses in CA3 (Weisskopf et al., 1994; Zalutsky and Nicoll, 1990), and subsequently for several other synapses in the CNS, e.g. cortical afferents to lateral amygdala and thalamus, cerebellar parallel fibers onto Purkinje cells, and many others (Yang and Calakos, 2013). The mechanisms of presynaptic LTP were studied most extensively at MF-CA3 synapses, formed between the axons of dentate gyrus granule cells and the dendrites of CA3 pyramidal cells (Nicoll and Schmitz, 2005). Here, high-frequency electrical stimulation of axons elicits first a strong post-tetanic potentiation of transmitter release, which decays within tens of minutes and is followed by LTP expressed as long-lasting increase of transmission. Alternatively, direct pharmacological activation of adenylyl cyclases by forskolin or application of the cAMP analogue Sp-cAMPS induces presynaptic potentiation (Huang et al., 1994; Tong et al., 1996; Weisskopf et al., 1994). However, both electrical and pharmacological inductions are not cell type-specific and difficult to implement in living animals. This lack of minimally invasive and cell-specific tools for controlling presynaptic short-and long-term potentiation is one of the reasons why despite decades of research on presynaptic plasticity, the molecular mechanisms downstream of cAMP are still not completely understood, nor the computational and behavioral relevance of presynaptic LTP (Monday et al., 2018). In order to fill this gap, here we establish synaptoPAC, an optogenetic tool for cell type-specific manipulation of presynaptic plasticity directly at the axonal terminal.

## Results

### SynaptoPAC design and functional validation

Optogenetic manipulations allow cell type-specific control of biochemical or electrical processes with high spatiotemporal precession, which can be further refined by subcellular targeting strategies (Rost et al., 2017). We reasoned that a presynaptically targeted light-activated adenylyl cyclase would enable optogenetic potentiation of transmitter release by a light-triggered increase of presynaptic cAMP. The small photoactivated adenylyl cyclase bPAC is ideally suited for this purpose because it is highly sensitive to blue light, shows low dark activity, exhibits a >100 fold increased cAMP-production in light, and is well expressed in neurons (Stierl et al., 2011). For the presynaptic targeting of bPAC we adopted a strategy previously established for optogenetic tools and fluorescent sensors (Granseth et al., 2006; Rost et al., 2015): By fusing bPAC together with a red fluorophore to the cytosolic C-terminus of the synaptic vesicle protein synaptophysin, we created synaptoPAC, a presynaptically targeted version of bPAC (Figure 1A). When coexpressed with the cAMP-sensitive K^+^-channel SthK (Brams et al., 2014) in ND7/23 cells, light-activation of synaptoPAC triggered outward currents similarly to untargeted bPAC, demonstrating efficient cAMP production by the enzyme (Figures 1B and 1C). In cultures of rat hippocampal neurons, costainings for the glutamatergic vesicle marker VGLUT1 revealed a punctate expression of synaptoPAC, largely overlapping with VGLUT1 (Figure 1D). Thus, fusion of bPAC to synaptophysin does not interfere with its enzymatic function, and facilitates presynaptic accumulation of the construct. Next, we recorded excitatory postsynaptic currents (EPSCs) from autaptic cultures of hippocampal granule cells, which show forskolin-induced potentiation of transmitter release (Rost et al., 2010; Tong et al., 1996). Granule cells were identified by their specific sensitivity of the transmitter release towards agonists of group II metabotropic glutamate receptors (Figure 2A, see Methods for details). After pharmacological characterization and establishing a stable baseline, we triggered synaptoPAC-mediated cAMP production with 12 1-s pulses of blue light at 0.2 Hz. In granule cells, photostimulation increased EPSCs to 129 ± 4% of baseline, but had no effect on transmission in non-granule cells (Figure 2B), indicating that the potentiation of synaptic transmission via photo-induced production of cAMP might be a cell-type-specific effect. Blue-light illumination occluded a further increase of EPSCs by forskolin in granule cells (Figure 2C). The optogenetic potentiation also altered the short-term plasticity of synaptic transmission, as it reduced the amplitude ratio of two EPSCs evoked at 25 Hz (Figure 2D). Such a decrease in the paired-pulse ratio (PPR) is indicative for a presynaptic origin of the effect, and often correlates with an increase in the release probability (Abbott and Regehr, 2004; Zucker and Regehr, 2002). In contrast, synaptoPAC activation did not alter the PPR in non-granule cells. Light-triggered elevation of presynaptic cAMP caused an increase in the frequency of miniature EPSCs (mEPSCs), but did not affect mEPSC amplitude, independent of the cell type (Figure S1). In summary, our experiments in neuronal cultures indicate that synaptoPAC acts via a presynaptic mechanism, and affects evoked release in a synapse-specific manner.

**Figure 1.**
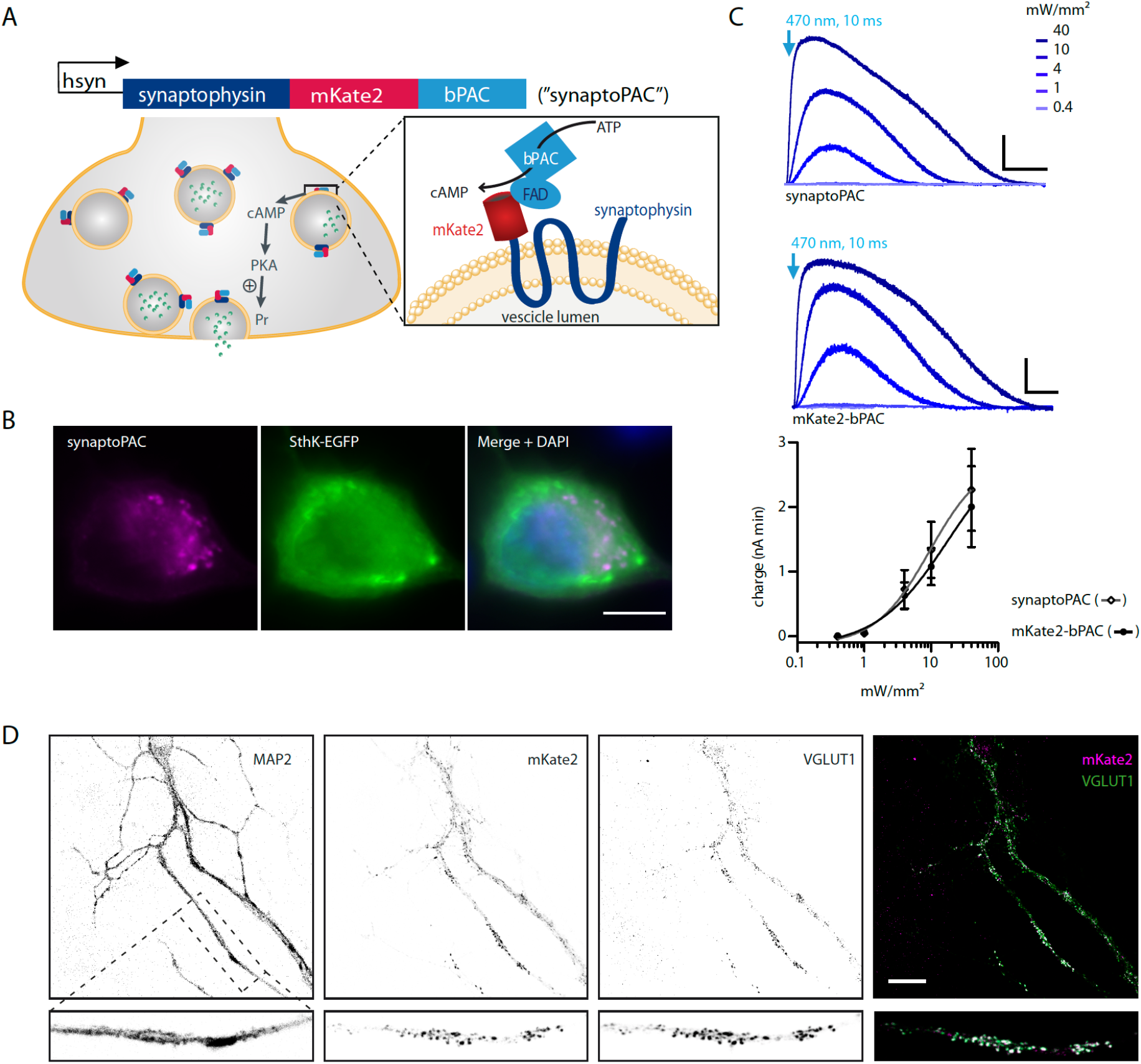
Design and validation of synaptoPAC. (**A**) Illustration of the presynaptic targeting strategy for bPAC, and the optogenetic induction of presynaptic plasticity by synaptoPAC activation. (**B**) Fluorescent microscope images of an ND7/23 cell expressing synaptoPAC and SthK-EGFP. Scale bar: 10 μm. (**C**) 10 ms flashes of 470 nm light triggered intensity-dependent outward currents in ND7/23 cells co-expressing the cAMP-gated SthK channel together with synaptoPAC (n = 8, N = 2) or untargeted bPAC (n = 8, N = 2). Scale bars: 200 pA, 20 s. (**D**) Fluorescent microscope images of cultured neurons and close up of dendrite showing colocalization (white in merge) of the mKate2 signal indicating synaptoPAC expression (magenta) and VGLUT1 as presynaptic marker (green). MAP2 was used as dendritic marker (not shown in merge). Scale bar: 20 μm.

**Figure 2.**
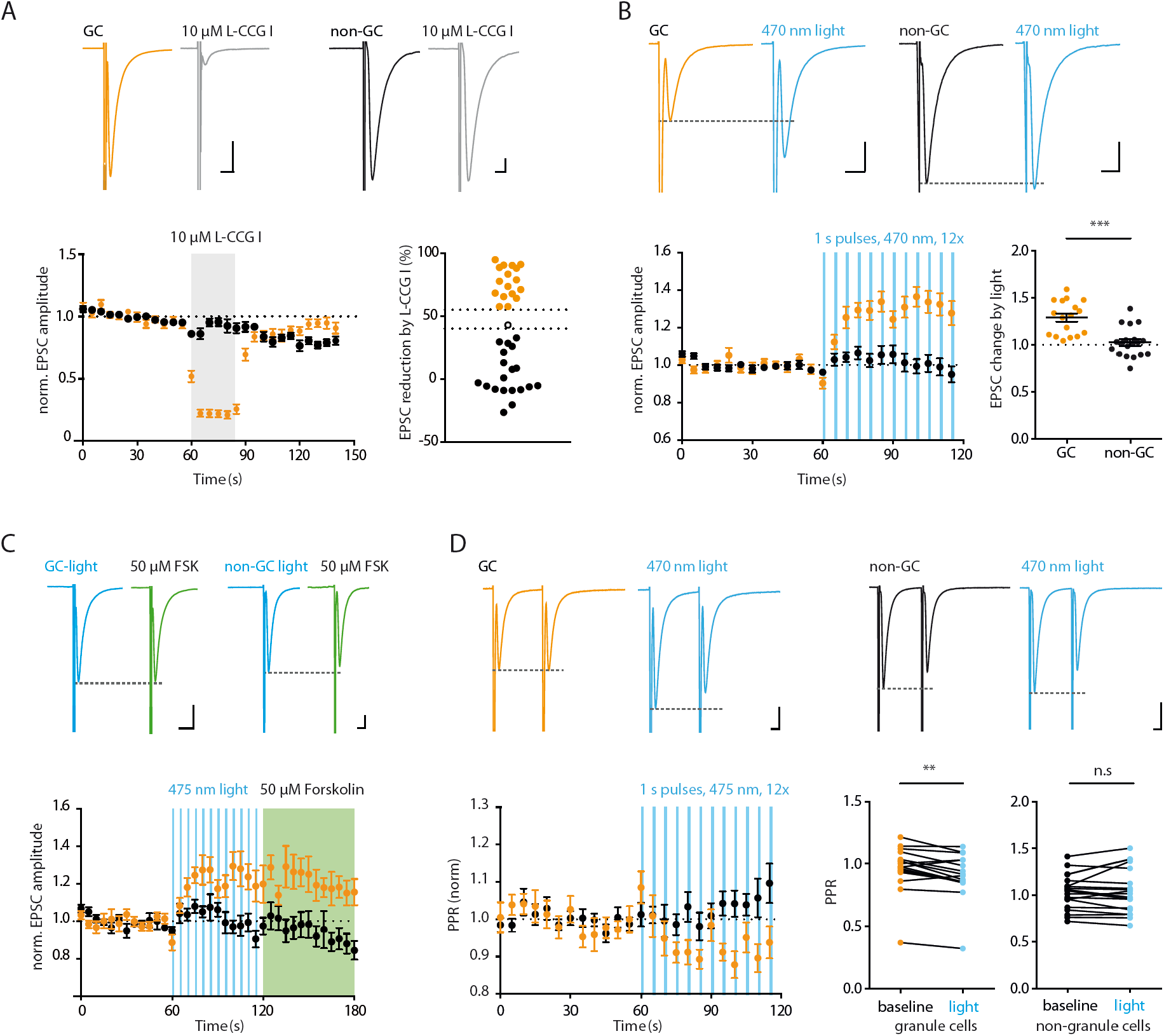
SynaptoPAC potentiates evoked synaptic transmission specifically in cultured granule cells. (**A**) Example traces illustrating the effect of the group II metabotropic glutamate receptor agonist L-CCG I (10 μM) on two types of hippocampal excitatory neurons in autaptic cultures. Scale bars, 10 ms, 1 nA. Neurons in which L-CCG I decreased the EPSC amplitude by >55% were classified as granule cells (GCs). Cells with <40% reduction of the EPSC amplitude by L-CCG I were defined as non-granule cells (non-GCs), others (open circle) were not included in further analysis. Time plot shows the reversible L-CCG I effect on EPSC amplitudes recorded from GCs (orange; n = 17, N = 4) and non-GCs (black; n = 19, N = 5). (**B**) EPSCs recorded before and after synaptoPAC activation from a GC and non-GC. Scale bars: 10 ms, 1 nA. Light pulses (470 nm, 70 mW/mm^2^, 12 × 1 s at 0.2 Hz) increased EPSC amplitudes in synaptoPAC-expressing GCs by 1.29 ± 0.04 (orange; n = 17, N = 4), but had no effect in synaptoPAC-expressing non-GCs (black; 1.03 ± 0.03, n = 19, N = 5; p < 0.0001; unpaired t-test). (**C)** Potentiation of EPSCs by activation of synaptoPAC (blue) and by subsequent application of 50 μM forskolin (FSK; green). Scale bars: 10 ms, 1 nA. In GCs, FSK increased the EPSC amplitude by 1.21 ± 0.07 (orange; n = 10, N = 4), not further increasing light induced potentiation. In non-GCs FSK did also not increase EPSCs (0.94 ± 0.06; black; n = 9, N = 3). (**D**) Paired EPSCs evoked at 25 Hz in a GC (orange) and a non-GC (black). Scale bars: 1 nA, 10 ms. Light stimulation significantly decreased the paired-pulse ratio in GCs (n = 17, N = 4; p = 0.002; Wilcoxon test) but had no effect on the paired-pulse ratio in non-GC (n = 19, N = 5; p = 0.5, paired t-test).

### SynaptoPAC potentiates transmission at MF-CA3 synapses

Next, we tested whether synaptoPAC was able to potentiate transmission at hippocampal MF-CA3 synapses, the classical preparation to study presynaptic forms of LTP. We injected AAVs encoding synaptoPAC into the dentate gyrus of wildtype mice *in vivo*. After three to four weeks, we prepared acute hippocampal slices, and found strong synaptoPAC expression in dentate gyrus granule cells and the MF tract (Figure 3A). We electrically stimulated MF transmission and recorded field excitatory postsynaptic potentials (fEPSPs) in area CA3. SynaptoPAC was activated by blue light pulses locally at MF terminals through a 40x objective in the area of the recording electrode (Figure 3B). MF-fEPSP amplitudes increased to 313 ± 28% within 5 min after the start of the optical induction protocol (Figure 3C). Subsequently, fEPSPs decreased, but were still potentiated by 126 ± 0.3% after 20 to 30 min post induction. At the end of each recording we verified the MF origin of the signal by applying the metabotropic glutamate receptor agonist DCG-IV (Figure 3C), which specifically suppresses release from MF, but not from neighboring associational-commissural fiber synapses in CA3 (Kamiya et al., 1996). Next, we examined whether synaptoPAC-induced plasticity was specific for synapses that express cAMP-dependent presynaptic LTP. For this we expressed synaptoPAC in CA3 pyramidal neurons (Figures 3D and 3E), and recorded fEPSPs from Schaffer collateral synapses in CA1, which display an NMDAR-dependent, postsynaptic expression mechanism of LTP (Herring and Nicoll, 2016). Optical activation of synaptoPAC with the same protocol that elicited strong potentiation in MF did not induce significant potentiation on Schaffer collateral synapses (Figures 3F and 3G), however, these synapses showed potentiation following tetanic electrical stimulation (Figure 3F). Together, the experiments in acute hippocampal slices confirm a synapse-specific effect of synaptoPAC.

**Figure 3.**
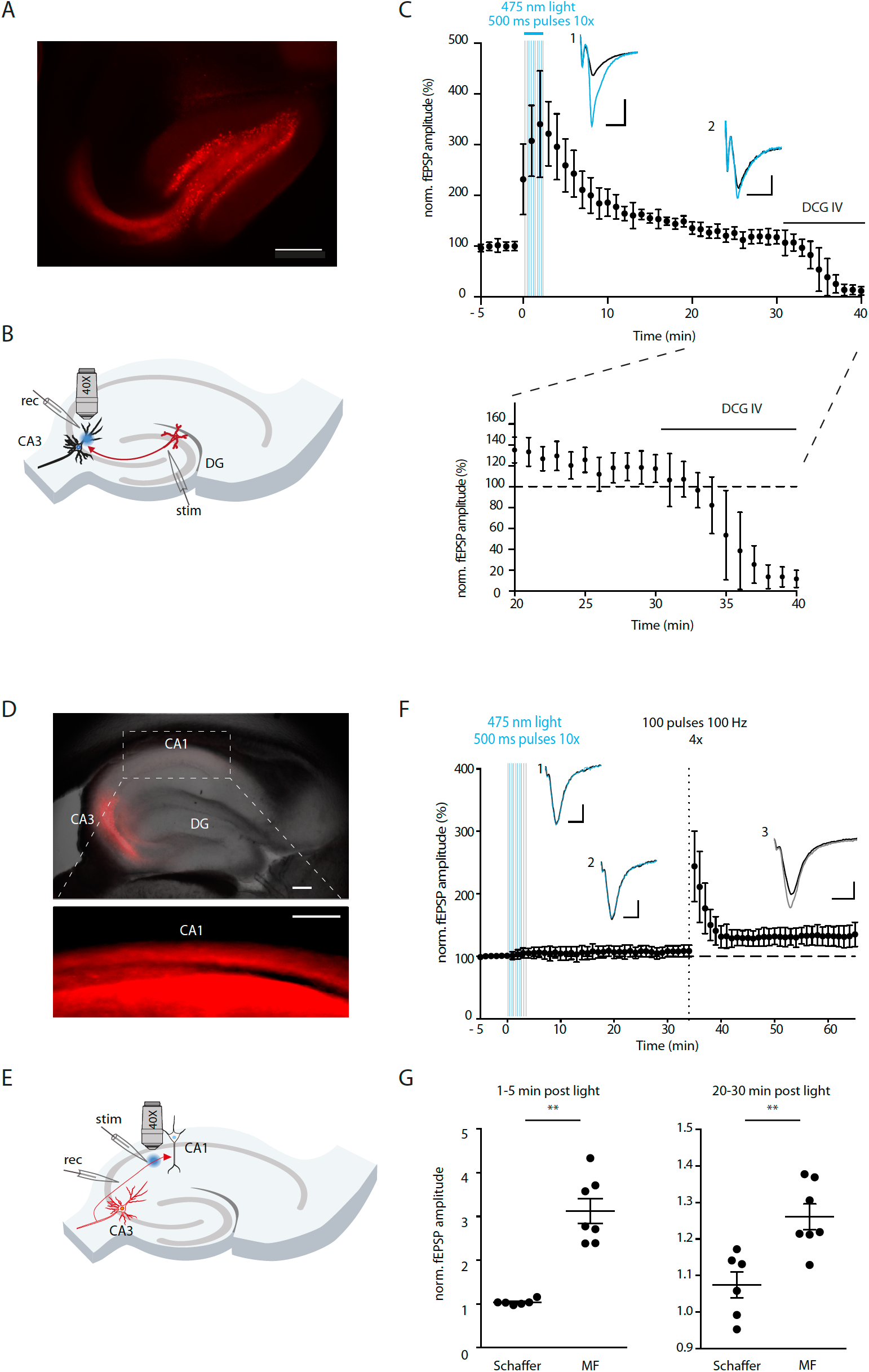
SynaptoPAC activation increases transmission at mossy fiber-CA3, but not at Schaffer collateral-CA1 synapses. **(A)** Fluorescent image of mScarlet confirming synaptoPAC expression in the dentate gyrus and mossy fibers. Scale bar, 200 μm. **(B)** Schematic of the recording configuration in acute hippocampal slices for MF-CA3 synapses, with optical stimulation in *stratum lucidum* in CA3. **(C)** Optical activation with 500 ms pulses of 470 nm light (11 mW/mm^2^ repeated 10 times at 0.05 Hz) acutely increased MF-fEPSPs by 3.13 ± 0.28, and triggered LTP of MF-fEPSP (1.26 ± 0.034 increase) lasting >20 min. Example traces showing baseline synaptic transmission (black), (1) potentiated transmission in the first 5 min after illumination (blue; scale bar, 200 μV, 20 ms), and (2) LTP 20-30 min after optical induction (scale bar: 100 μV, 20 ms). Only experiments that showed >80% reduction of fEPSP amplitudes by application of DCG IV (1 μM) were considered as mossy fiber recordings (n = 7 slices, 6 mice). **(D)** Overlay of DIC and fluorescent image showing synaptoPAC expression in area CA3 and in Schaffer collaterals (inset). Scale bars: 200 μm. **(E)** Scheme of the electrophysiological recording configuration at Schaffer collateral-CA1 synapses. **(F)** In Schaffer collateral synapses expressing synaptoPAC, blue light stimulation had no immediate effect in the first 5 min post illumination (1) (1.04 ± 0.02), nor did it cause strong LTP after 20-30 min (2) (1.07± 0.03). However, these synapses showed potentiation (3) (1.32 ± 0.06) following 4 trains of 100 stimuli at 100 Hz (n = 6 slices, 3 mice. Scale bars: 100 μV, 10 ms. **(G)** Direct comparison of the synaptoPAC effect on transmission at MF-CA3 synapses and CA3-CA1 synapses. In both cases the effect size induced by synaptoPAC is significantly higher in mossy fibers synapses compared to Schaffer collaterals synapses (1-5 min: p = 0.0006; 20-30 min: p = 0.02, both Mann Whitney test).

### SynaptoPAC activation alters presynaptic release probability

One of the distinguishing characteristics of MF-CA3 synapses is the low basal release probability, which results in pronounced short-term facilitation of synaptic transmission upon repetitive activation (Nicoll and Schmitz, 2005). LTP in MF-CA3 synapses increases transmitter release probability, which reduces the dynamic range of the short-term plasticity of release (Gundlfinger et al., 2007; Weisskopf et al., 1994; Xiang et al., 1994; Zalutsky and Nicoll, 1990). To test whether optogenetic potentiation similarly alters MF release properties, we stimulated MF-fEPSPs with 5 pulses at 25 Hz and 20 pulses at 1 Hz, both before and after blue-light stimulation (Figure 4A). Indeed, illumination significantly decreased both the PPR of the second to the first and of the fifth to the first fEPSP (Figure 4B). Optogenetic potentiation also caused a significant decrease in 1 Hz frequency facilitation (Figure 4C). Both test protocols indicate that synaptoPAC-driven MF potentiation changes the release probability. Of note, the facilitation of MF-fEPSPs during 1 Hz stimulation in slices expressing synaptoPAC was 440% before blue light exposure, similar to published values of 400 to 600% recorded under comparable conditions (Kukley et al., 2005; Moore et al., 2003; Mori-Kawakami et al., 2003), which indicates that expression of synaptoPAC did not *per se* alter release properties. Taken together, synaptoPAC enables light-controlled production of cAMP at presynaptic terminals, which increases transmitter release specifically in synapses sensitive to cAMP.

**Figure 4.**
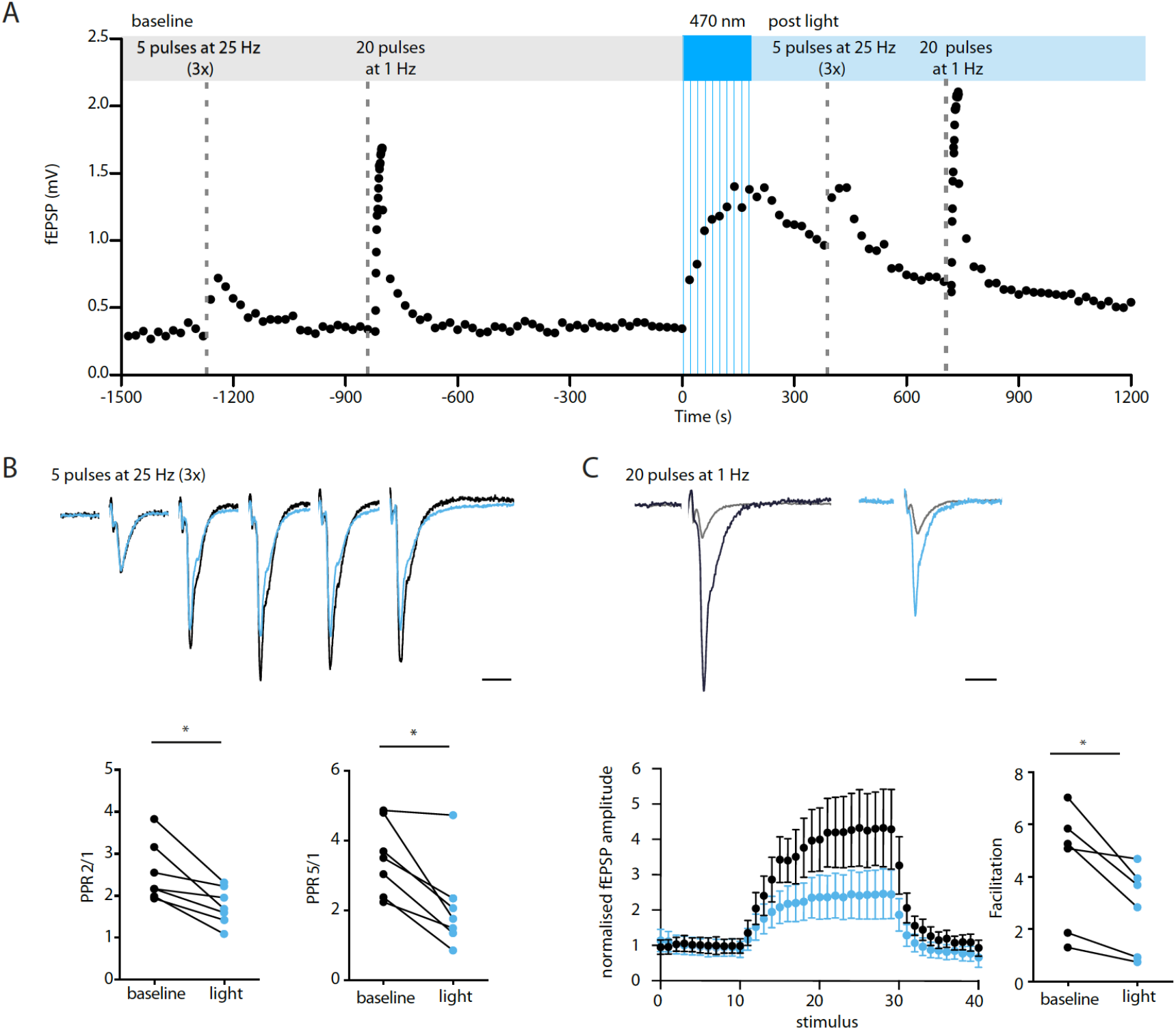
SynaptoPAC mediates potentiation by a presynaptic mechanism. (**A**) Typical representation of the recording protocol for MF short-term plasticity: during baseline MF were stimulated electrically with 5 pulses at 25 Hz and with 20 pulses at 1 Hz. Afterwards, synaptoPAC was activated by 10 pulses of 470 nm light (500 ms, 11 mW/mm^2^, 0.2 Hz). The 25 Hz and 1 Hz stimulus trains were applied again 5 and 10 min after optical induction, respectively. For the 25 Hz trains, only the first fEPSP amplitude was plotted. **(B)** Representative traces of paired pulse MF-fEPSP before (black) and after light stimulation (blue), scaled to the amplitude of the first fEPSP. Scale bar: 20 ms. SynaptoPAC induced potentiation significantly decreased the paired-pulse ratio between the second and the first fEPSPs (n =7 slices, 6 mice; p = 0.02, Wilcoxon signed rank test), and the fifth and first fEPSPs (p = 0.02, Wilcoxon signed rank test). **(C)** Traces showing the first and last MF-fEPSP evoked by a train of 20 stimuli applied at 1 Hz before (black) and after (blue) blue-light illumination. Traces normalized to the first fEPSP in baseline. Scale bar: 20 ms. 1 Hz facilitation significantly decreased after synaptoPAC induced potentiation (n = 7 slices, 6 mice; p = 0.03, Wilcoxon signed rank test).

## Discussion

Tools enabling selective and remote-controlled induction of presynaptic plasticity have been requested to elucidate the mechanisms and functional relevance of presynaptic LTP (Monday et al., 2018). SynaptoPAC is well suited for such investigations: First, it is easily expressed via AAVs due to its small size, and allows cell type-specific expression via Cre-lox recombination and specific promoters. Second, bPAC is highly light sensitive, and its selective activation by blue light (Stierl et al., 2011) enables experimental combinations with optogenetic actuators and fluorescent sensors sensitive to light >520 nm (Bernal Sierra et al., 2018). Third, local illumination of axonal terminals is sufficient to trigger potentiation, allowing target area-specific manipulations. Finally, the effect of synaptoPAC activation on MF-fEPSPs was probably underestimated in our slice experiments, since in the virally transduced tissue both synaptoPAC-expressing and non-transduced cells contribute to the electrically evoked fEPSP. Interestingly, the time course of synaptoPAC mediated potentiation of MF transmission in hippocampal slices matches the time-course of potentiation induced by tetanic-stimulation, with a strong initial increase that decays with a tau of approx. 20 min. This suggest that a transient increase of presynaptic cAMP is sufficient for post-tetanic potentiation at MF, providing an alternative mechanistic explanation for the enhanced transmission following bursts of activity (Regehr, 2012).

Our experiments in acute hippocampal slices on CA3-CA1 Schaffer collateral synapses expressing synaptoPAC revealed that a selective increase of presynaptic cAMP in these synapses is not sufficient to potentiate action potential-evoked transmission (Figure 3). Previously, forskolin in combination with phosphodiesterase inhibitors was used to increase transmission at Schaffer collateral synapses (Chavez-Noriega and Stevens, 1992, 1994; Frerking et al., 2001; Otmakhov et al., 2004). However, it is unclear whether this form of chemical LTP is purely presynaptic, if physiological activity can recapitulate the pharmacological stimulation, and to which extend cAMP-independent effects of forskolin contribute to it (Laurenza et al., 1989). In our hands, a selective increase of presynaptic cAMP by optogenetic means without pharmacological inhibition of phosphodiesterases did not induce strong potentiation at Schaffer collateral synapses nor in cultures of hippocampal non-granule cells. Our findings suggest that Schaffer collateral synapses lack key features of the molecular machinery required for presynaptic long-term plasticity at mossy fiber terminals, resulting in a synapse-specific effect of cAMP on evoked transmission.

Of note, and in line with previous reports (Maximov et al., 2007), synaptoPAC-driven elevation of presynaptic cAMP increased mEPSC frequency, irrespective of the glutamatergic cell type. Presynaptic cAMP promotes synapsin phosphorylation by PKA, causing dissociation of synapsin from vesicles, and spatial reorganization of vesicles in the terminal (Menegon et al., 2006; Patzke et al., 2019; Vaden et al., 2019). In future, optogenetic control of presynaptic cAMP by synaptoPAC will provide novel avenues to study such kinds of vesicle pool regulation.

SynaptoPAC-driven enhancement of release will be useful to study computational aspects of presynaptic potentiation, which alters short-term synaptic filtering dynamics by shifting spike transmission from a high-pass towards a low-pass mode (Abbott and Regehr, 2004). How this altered synaptic computation affects encoding, storage and recall of associative memories is mostly unknown (Rebola et al., 2017). So far, causative behavioral studies relied on either genetic ablation of major presynaptic molecules, or drugs of abuse acting on presynaptic metabotropic receptors such as cannabinoids, opioids, and cocaine (for review see (Kauer and Malenka, 2007; Monday et al., 2018)). However, since these manipulations are slow and often systemically, it is difficult to clearly relate synaptic alterations and behavioral consequences. Presynaptic plasticity has also been associated with pathophysiological brain conditions such as schizophrenia, autism spectrum disorders, and addiction, but it remains unclear whether aberrant presynaptic plasticity contributes causally or results from the disease (Monday and Castillo, 2017). We expect that synaptoPAC represents a versatile optogenetic tool to unravel some of the outstanding questions of presynaptic plasticity, both under physiological and pathophysiological conditions.

## Acknowledgements

We thank Katja Pötschke, Bettina Brokowski, Anke Schönherr, Susanne Rieckmann and Katja Czieselsky for excellent technical assistance. This work was supported by grants from the Deutsche Forschungsgemeinschaft (DFG; German Research Foundation) under Germany’s Excellence Strategy – EXC-2049 – 390688087 to D.S. and C.R., SPP1926 (B.R.R), SPP 1665 (D.S.), SFB 1315 (D.S.), SFB 958 (C.R., D.S.). The plasmid encoding bPAC was kindly provided by the laboratory of Peter Hegemann, Humboldt University Berlin, Germany. SthK-EGFP was kindly provided by Franziska Schneider-Warme, Institute for Experimental Cardiovascular Medicine, University of Freiburg, Germany.

## Author contributions

S.O., L.M-V., A.S. and B.R.R performed the experiments. S.O. and B.R.R. analyzed the data. B.R.R. designed the tool, and B.R.R., D.S. and C.R. conceptualized the research. B.R.R and D.S. acquired funding for the project and supervised the work. B.R.R and S.O. wrote the paper with input from all authors. All authors approved submission of this manuscript.

---

**Figure S1.**
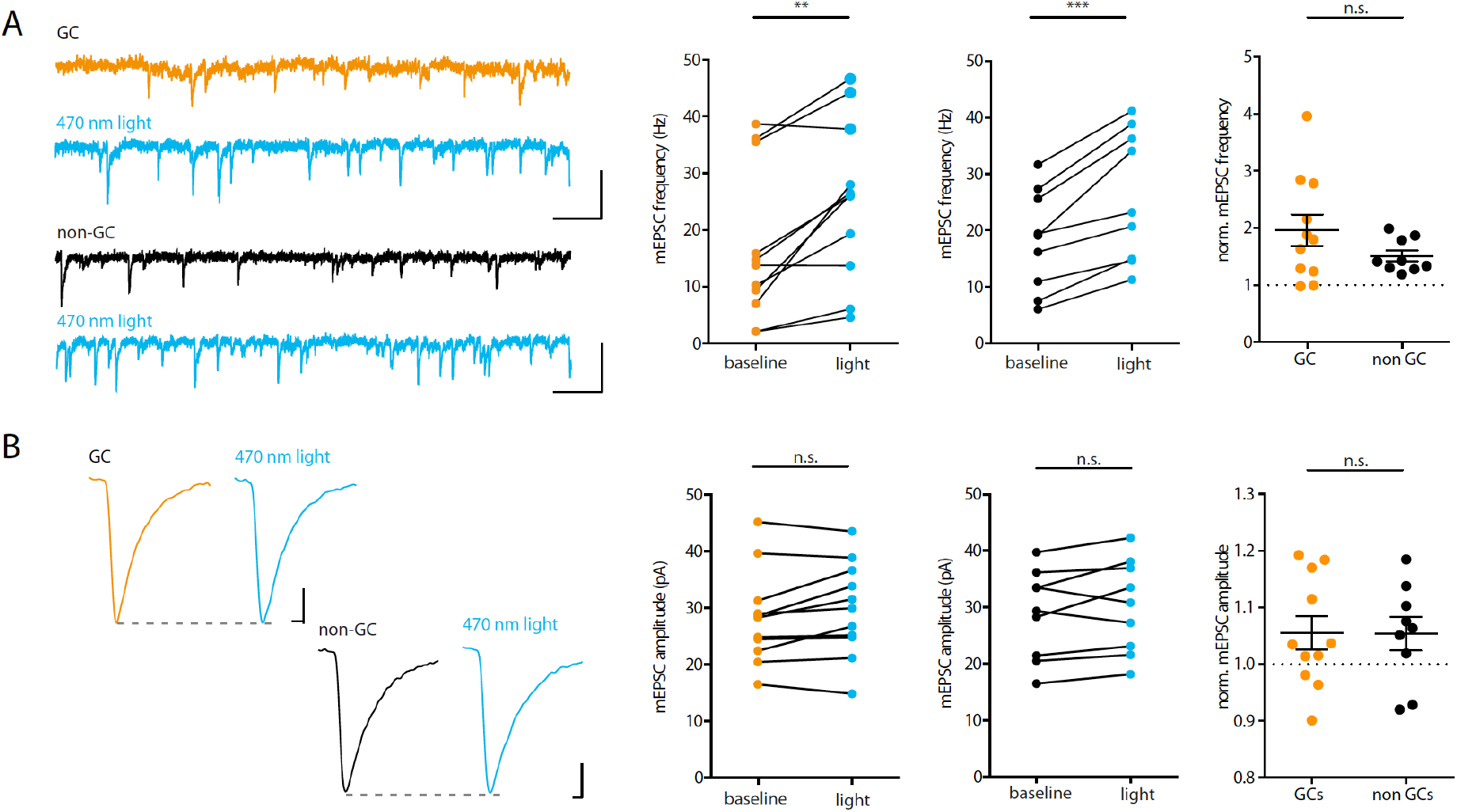
SynaptoPAC activation increases mEPSCs frequency but not amplitude. **(A)** Example traces of mEPSCs of a GC (orange) and a non-GC (black) before and after light (blue). Scale bar: 50 pA, 100 ms. Light exposure increased the frequency of mEPSCs significantly compared to baseline both in GCs (baseline: 16.89 ± 4.09, light: 25.37 ± 4.21; n = 11, N = 5; p = 0.002, paired t-test) and in non-GCs (baseline: 18.2 ± 2.98, light: 26.2 ± 3.85; n = 9, N = 4; p = 0.0003, paired t-test). The mEPSC frequency change in GCs was not significantly different from that in non-GCs (p = 0.13, unpaired t-test). **(B)** Example traces of averaged mEPSCs of a GC (orange) and of a non-GC (black) before and after light (blue). Scale bars: 5 pA, 1 ms. SynaptoPAC activation did not affect mEPSC amplitude in GCs (baseline: 28.3 ± 2.49, light: 29.8 ± 2.50; n = 11, N = 5; p = 0.082, paired t-test) nor in non-GCs (baseline: 28.8 ± 2.62, light: 30.2 ± 2.74; n = 9, N = 4; p = 0.148, paired t-test). The mEPSC amplitudes of GCs and non-GCs post light normalized to before light were not significantly different (p = 0.97, unpaired t-test).

## Materials and Methods

### Animal experiments

All animal experiments were carried out according to the guidelines stated in Directive 2010/63/EU of the European Parliament on the protection of animals used for scientific purposes and were approved by the local authorities in Berlin (Berlin state government/Landesamt für Gesundheit und Soziales, license number G0092/15, T0100/03 and T0073/04), Germany.

### Molecular biology

For initial tests in neuronal cell cultures, we fused the coding sequence of mKate2 and bPAC (from the soil bacterium *Beggiatoa*) to the 3’ end of the rat synaptophysin coding sequence by Gibson assembly. The resulting fusion construct (synaptophysin-mKate2-bPAC) was then transferred into a lentiviral expression vector via PCR, which allowed expression under control of the neuron-specific human synapsin (hSyn) promoter. This construct was used for the characterization in neuronal cultures (Figures 1 and 2). Since mKate2-fluorescence was very dim and difficult to detect in live-cell imaging, we replaced the fluorophore by mCherry, and transferred the construct into an adeno-associated virus (AAV)-expression vector for more efficient expression of synaptoPAC *in vivo*. In a final step of optimization, we replaced mCherry by mScarlet, since mCherry caused unintended protein aggregations in neurons (data not shown). All viruses (lentivirus and AAV serotype 9) were provided by the Viral Core Facility of the Charité Berlin.

### Autaptic granule cell culture

Autaptic cultures of dentate gyrus granule cells were prepared as described previously (Rost et al. 2010). In brief, dentate gyri of P0-P1 Wistar rats of either sex were dissected and separated from the rest of the hippocampus. Cells were subsequently plated on islands of glial cells at a density of 1.25 × 10^4^ cells/cm^2^ in 6-well plates in Neurobasal-A supplemented with 2% B27 and 0.2% penicillin/streptomycin (Invitrogen/ Thermo Fisher, Waltham, MA). Lentiviral particles encoding synaptoPAC were added 1–4 days after plating. Cells were used for electrophysiological recordings after 15–21 days *in vitro*.

### Cell culture of ND7/23 cells

ND7/23 cells (ECACC 92090903, Sigma-Aldrich/ Merck, Darmstadt, Germany) were cultured in Dulbecco’s Modified Eagle’s Medium (DMEM) supplemented with 10% fetal calf serum and 0.1% penicillin/streptomycin (Invitrogen) at 37°C, 5% CO_2_. 24 h prior to transfection, cells were seeded onto 15 mm coverslips coated with collagen and poly-D-lysin. Cells were transfected using X-tremeGENE 9 (ratio 3:1 μl/μg DNA) according to the manufacturer’s protocol (Sigma-Aldrich). DNA encoding PAC constructs and SthK-EGFP was used at a 3:1 ratio. Two days after transfection we performed whole cell voltage clamp recordings on transfected cells identified by red fluorescence.

### Immunocytochemistry

At DIV 15–21, cultured neurons were rinsed with PBS and fixed with 4% PFA in PBS for 10 min, then washed 3 times in PBS. Samples were permeabilized with 0.5% Triton-X100 for 5 min and blocked for 30 min in 2% normal goat serum in 0.1% Triton-X100. Samples were incubated with rabbit anti-RFP (against mKate2; 1:200; Invitrogen), chicken anti-microtubule-associated protein 2 (MAP2; 1:2000; Chemicon/Merck), and guinea pig anti-vesicular glutamate transporter 1 (VGLUT1; 1:4000; Synaptic Systems, Göttingen, Germany) primary antibodies at room temperature for 1.5 h. Coverslips were washed 3 times with PBS before incubation with secondary antibodies conjugated with AlexaFluor-488, −555, or −647 (1:500; Invitrogen) at room temperature for 1 h. Finally, coverslips were washed 3 times with PBS and mounted with Mowiol (Carl Roth, Karlsruhe, Germany).

### Confocal microscopy

Neurons were imaged on an upright TCS SP5 confocal microscope (Leica Microsystems, Wetzlar, Germany). The following laser lines were used to illuminate fluorescent specimens through a 63x oil immersion objective (1.4 NA): 488 nm (Argon laser), 568 nm (solid state), and 633 nm (Helium, Neon). Images were acquired at 1024 × 1024 pixels.

### Electrophysiology on ND7/23 cells and neuronal cultures

Whole cell voltage clamp recordings were performed on an IX73 inverted microscope (Olympus, Shinjuku, Tokyo, Japan) using a Multiclamp 700B amplifier under the control of a Digidata 1550 AD board and Clampex 10 software (all Molecular Devices, San José, CA). All recordings were performed at room temperature. Data was acquired at 10 kHz and filtered at 3 kHz. A TTL-controlled LED system (pE4000, CoolLED, Andover, UK) was coupled into the back port of the IX73 microscope by a single liquid light guide. Fluorescent light was passed through a quadband filter set (F66-415, AHF, Tübingen, Germany) and an Olympus UPLSAPO 20×, NA 0.75 objective. For visualization of mKate2 fluorescence, we used a triband filter set (AHF F66-502) with 575/25 nm excitation filter. Extracellular solution contained (in mM): 140 NaCl, 2.4 KCl, 10 HEPES, 10 glucose, 2 CaCl_2_, and 4 MgCl_2_ (pH adjusted to 7.3 with NaOH, 300 mOsm). The intracellular solution contained (in mM): 135 K-gluconate, 17.8 HEPES, 1 EGTA, 4.6 MgCl_2_, 4 Na_2_-ATP, 12 disodium creatine phosphate, and 50 U/ml creatine phosphokinase, pH adjusted to 7.3 with KOH, 300 mOsm. Unless otherwise stated, all chemicals were purchased from Tocris, Merck or Carl Roth. ND7/23 cells co-expressing SthK-EGFP and synaptoPAC or mKate2-bPAC were voltage clamped at −60 mV. Photoactivated adenylyl cyclases were stimulated by 10 ms flashes of 470 nm light with varying intensity, which elicited cAMP-triggered K^+^ currents.

In whole-cell voltage clamp recordings from cultured neurons, membrane potential was set to −70 mV. Paired EPSCs were evoked every 5 s by triggering two unclamped action potentials at 25 Hz using 1 ms-depolarizations of the soma to 0 mV. Granule cells were distinguished from other glutamatergic hippocampal neurons by the sensitivity of synaptic transmission to application of the group II mGluR agonist L-CCG I ((2S,1’S,2’S)-2-(Carboxycyclopropyl)glycines; 10 μM; Tocris, Bristol, UK). We classified neurons as granule cells if L-CCG I reduced their transmission by >55%, neurons which showed <40% reduction of the EPSC were classified as non-granule cells. SynaptoPAC was activated by 12 flashes (1s) of 470-nm light at 0.2 Hz at 70 mW/mm^2^. The PPR was calculated as the ratio from the second and first EPSC amplitude with an interstimulus interval of 40 ms. Data was analyzed using AxoGraph X (AxoGraph, Sydney, Australia). EPSC potentiation and PPR changes were calculated from averaged EPSCs during baseline and light stimulation. For detection of miniature EPSCs, we filtered the recordings post hoc with a digital 1-kHz low-pass filter, and applied a template-based algorithm implemented in AxoGraph X. Frequency and amplitude of mEPSCs were corrected for false positives events, which we estimated by running the event detection with an inverted template (Rost et al., 2015). Recordings were excluded from the mEPSC analysis if false positive events occurred at a frequency of >1Hz.

### Stereotactic surgeries

Wild type C57/BL6-N mice (P25–P31) were injected with 200 nl of AAV9.hSyn:synaptophysin-mScarlet-bPAC using a glass micropipette attached to a 10-μL Hamilton microsyringe. Mice were anesthetized with 1.5% isoflurane during surgery. Injections were aimed at the following coordinates relative to bregma: anteroposterior (AP): −1.93 mm, mediolateral (ML): +1.5 mm, dorsoventral (DV): −2.0 mm for the DG; AP: −1.75 mm, ML: 2.00 mm, DV: −2.1 mm for CA3. All animals were allowed to recover for at least 3 weeks before slice preparations and recording.

### Electrophysiology on hippocampal slices

Animals were anaesthetized under isoflurane and decapitated. Under continuous safe light illumination, brains were quickly removed and placed in ice-cold sucrose-based artificial cerebrospinal fluid (ACSF) containing (in mM): 87 NaCl, 26 NaHCO_3_, 10 glucose, 2.5 KCl, 3.5 MgCl_2_, 1.25 NaH_2_PO_4_, 0.5 CaCl_2_, and 75 sucrose, saturated with 95% O_2_ and 5% CO_2_. Tissue blocks containing the hippocampal formation were mounted on a vibratome (Leica VT 1200S, Leica Microsystems). Sagittal slices were cut at 300-μm thickness and incubated in sucrose-based ACSF at 35 °C for 30 min. Before the recording, slices were transferred into a submerged chamber filled with recording ACSF containing the following (in mM): 119 NaCl, 26 NaHCO_3_, 10 glucose, 2.5 KCl, 1.3 MgCl_2_, 1 NaH_2_PO_4_, and 2.5 CaCl_2_, saturated with 95% O_2_ and 5% CO_2_. Slices were stored in ACSF for 0.5–6 h at room temperature.

MF-fEPSPs recordings were performed at 21–24°C in a submerged recording chamber perfused with ACSF. MF were stimulated at 0.05 Hz using a low-resistance glass electrode filled with ACSF, which was placed close to the internal side of granule cell layer of the dentate gyrus. The recording electrode was placed in *stratum lucidum* of area CA3. The MF origin of fEPSPs was verified by the pronounced facilitation of synaptic response amplitudes upon 1 Hz stimulation or paired pulse stimulation at 25 Hz. In addition, we confirmed that fEPSPs were generated specifically by MF by application of the type II metabotropic glutamate receptor agonist DCG IV ((2S,2’R,3’R)-2-(2’,3’-dicarboxycyclopropyl)glycine; 1 μM) at the end of each recording. DCG IV specifically suppresses transmitter release from MF terminals in CA3, but not from neighboring associational-commissural fiber synapses (Kamiya et al., 1996). Only responses that were inhibited by >80% were accepted as mossy fiber signals.

Recordings from Schaffer collateral synapses in CA1 were performed at 21–24°C in a submerged recording chamber perfused with ACSF at 2 ml/min. The stimulation electrode was placed in *stratum radiatum* of CA3 and the recording electrode in the *stratum radiatum* of CA1. Field EPSPs were stimulated at 0.05 Hz. A stable baseline of fEPSP amplitudes was recorded for at least 10 minutes before optical stimulation. In CA1 fEPSP recordings, we verified the fidelity of synaptic plasticity at the end of each recording (>30 min post optical stimulation) by electrically inducing LTP by four high frequency trains (100 pulses at 100 Hz, 10 s inter train interval). Only slices that displayed >10% increase of the fEPSP 20-30 min after tetanic stimulation and showed <15% change in presynaptic fiber volley over the course of the whole recording were included in the analysis.

In both type of recordings, optical potentiation was induced by 10 pulses of blue light at 0.05 Hz from a 470 nm LED (pE300, CoolLED), which was passed through a 474/23 nm filter (AHF F39-474). Light was applied via an Olympus LUMPLFLN 40x (NA 0.8) water immersion objective, which was placed in *stratum lucidum* of CA3 for MF recordings and in *stratum radiatum* of CA1 for Schaffer collateral recordings. Data was acquired using a Multiclamp 700B amplifier, Digidata 1550A AD board, and Clampex 10.0 recording software (all Molecular Devices). In some recordings, data was acquired using a Multiclamp 700B amplifier, a BNC-2090 interface board with PCI 6035E A/D board (National Instruments, Austin, TX) and IGOR Pro 6.12 as recording software (WaveMetrics, Lake Oswego, OR). Signals were sampled at 10 kHz and filtered at 3 kHz. Recordings were analyzed using Clampfit (Molecular Devices), IGOR Pro, or AxographX.

### Statistics

No randomization or blinding was performed in the study. We did not perform a power analysis to determine sample size prior to the experiments, since the aim of our study was to establish a new technology without prior knowledge on effect size and variability. Data are presented as mean ± SEM.

## References

Abbott, L.F., and Regehr, W.G. (2004). Synaptic computation. Nature 431, 796–803.

Bernal Sierra, Y.A., Rost, B.R., Pofahl, M., Fernandes, A.M., Kopton, R.A., Moser, S., Holtkamp, D., Masala, N., Beed, P., Tukker, J.J., et al. (2018). Potassium channel-based optogenetic silencing. Nat Commun 9, 4611.

Brams, M., Kusch, J., Spurny, R., Benndorf, K., and Ulens, C. (2014). Family of prokaryote cyclic nucleotide-modulated ion channels. Proc Natl Acad Sci U S A 111, 7855–7860.

Chavez-Noriega, L.E., and Stevens, C.F. (1992). Modulation of synaptic efficacy in field CA1 of the rat hippocampus by forskolin. Brain Res 574, 85–92.

Chavez-Noriega, L.E., and Stevens, C.F. (1994). Increased transmitter release at excitatory synapses produced by direct activation of adenylate cyclase in rat hippocampal slices. J Neurosci 14, 310–317.

Ferguson, G.D., and Storm, D.R. (2004). Why calcium-stimulated adenylyl cyclases? Physiology (Bethesda) 19, 271–276.

Frerking, M., Schmitz, D., Zhou, Q., Johansen, J., and Nicoll, R.A. (2001). Kainate receptors depress excitatory synaptic transmission at CA3-->CA1 synapses in the hippocampus via a direct presynaptic action. J Neurosci 21, 2958–2966.

Granseth, B., Odermatt, B., Royle, S.J., and Lagnado, L. (2006). Clathrin-mediated endocytosis is the dominant mechanism of vesicle retrieval at hippocampal synapses. Neuron 51, 773–786.

Gundlfinger, A., Leibold, C., Gebert, K., Moisel, M., Schmitz, D., and Kempter, R. (2007). Differential modulation of short-term synaptic dynamics by long-term potentiation at mouse hippocampal mossy fibre synapses. J Physiol 585, 853–865.

Herring, B.E., and Nicoll, R.A. (2016). Long-Term Potentiation: From CaMKII to AMPA Receptor Trafficking. Annu Rev Physiol 78, 351–365.

Huang, Y.Y., Li, X.C., and Kandel, E.R. (1994). cAMP contributes to mossy fiber LTP by initiating both a covalently mediated early phase and macromolecular synthesis-dependent late phase. Cell 79, 69–79.

Kamiya, H., Shinozaki, H., and Yamamoto, C. (1996). Activation of metabotropic glutamate receptor type 2/3 suppresses transmission at rat hippocampal mossy fibre synapses. J Physiol 493 (Pt 2), 447–455.

Kauer, J.A., and Malenka, R.C. (2007). Synaptic plasticity and addiction. Nat Rev Neurosci 8, 844–858.

Kukley, M., Schwan, M., Fredholm, B.B., and Dietrich, D. (2005). The role of extracellular adenosine in regulating mossy fiber synaptic plasticity. J Neurosci 25, 2832–2837.

Laurenza, A., Sutkowski, E.M., and Seamon, K.B. (1989). Forskolin: a specific stimulator of adenylyl cyclase or a diterpene with multiple sites of action? Trends Pharmacol Sci 10, 442–447.

Maximov, A., Shin, O.H., Liu, X., and Sudhof, T.C. (2007). Synaptotagmin–12, a synaptic vesicle phosphoprotein that modulates spontaneous neurotransmitter release. J Cell Biol 176, 113–124.

Menegon, A., Bonanomi, D., Albertinazzi, C., Lotti, F., Ferrari, G., Kao, H.T., Benfenati, F., Baldelli, P., and Valtorta, F. (2006). Protein kinase A-mediated synapsin I phosphorylation is a central modulator of Ca2+–dependent synaptic activity. J Neurosci 26, 11670–11681.

Monday, H.R., and Castillo, P.E. (2017). Closing the gap: long-term presynaptic plasticity in brain function and disease. Curr Opin Neurobiol 45, 106–112.

Monday, H.R., Younts, T.J., and Castillo, P.E. (2018). Long-Term Plasticity of Neurotransmitter Release: Emerging Mechanisms and Contributions to Brain Function and Disease. Annu Rev Neurosci 41, 299–322.

Moore, K.A., Nicoll, R.A., and Schmitz, D. (2003). Adenosine gates synaptic plasticity at hippocampal mossy fiber synapses. Proc Natl Acad Sci U S A 100, 14397–14402.

Mori-Kawakami, F., Kobayashi, K., and Takahashi, T. (2003). Developmental decrease in synaptic facilitation at the mouse hippocampal mossy fibre synapse. J Physiol 553, 37–48.

Nicoll, R.A., and Schmitz, D. (2005). Synaptic plasticity at hippocampal mossy fibre synapses. Nat Rev Neurosci 6, 863–876.

Otmakhov, N., Khibnik, L., Otmakhova, N., Carpenter, S., Riahi, S., Asrican, B., and Lisman, J. (2004). Forskolin-induced LTP in the CA1 hippocampal region is NMDA receptor dependent. J Neurophysiol 91, 1955–1962.

Patzke, C., Brockmann, M.M., Dai, J., Gan, K.J., Grauel, M.K., Fenske, P., Liu, Y., Acuna, C., Rosenmund, C., and Sudhof, T.C. (2019). Neuromodulator Signaling Bidirectionally Controls Vesicle Numbers in Human Synapses. Cell 179, 498–513 e422.

Rebola, N., Carta, M., and Mulle, C. (2017). Operation and plasticity of hippocampal CA3 circuits: implications for memory encoding. Nat Rev Neurosci 18, 208–220.

Regehr, W.G. (2012). Short-term presynaptic plasticity. Cold Spring Harb Perspect Biol 4, a005702.

Rost, B.R., Breustedt, J., Schoenherr, A., Grosse, G., Ahnert-Hilger, G., and Schmitz, D. (2010). Autaptic cultures of single hippocampal granule cells of mice and rats. Eur J Neurosci 32, 939–947.

Rost, B.R., Schneider, F., Grauel, M.K., Wozny, C.C, G.B., Blessing, A., Rosenmund, T., Jentsch, T.J., Schmitz, D., Hegemann P., et al. (2015). Optogenetic acidification of synaptic vesicles and lysosomes. Nat Neurosci 18, 1845–1852.

Rost, B.R., Schneider-Warme, F., Schmitz, D., and Hegemann, P. (2017). Optogenetic Tools for Subcellular Applications in Neuroscience. Neuron 96, 572–603.

Stierl, M., Stumpf, P., Udwari, D., Gueta, R., Hagedorn, R., Losi, A., Gärtner, W., Petereit, L., Efetova, M., Schwärzel, M., et al. (2011). Light modulation of cellular cAMP by a small bacterial photoactivated adenylyl cyclase, bPAC, of the soil bacterium Beggiatoa. J Biol Chem 286, 1181–1188.

Takeuchi, T., Duszkiewicz, A.J., and Morris, R.G. (2014). The synaptic plasticity and memory hypothesis: encoding, storage and persistence. Philos Trans R Soc Lond B Biol Sci 369, 20130288.

Tong, G., Malenka, R.C., and Nicoll, R.A. (1996). Long-term potentiation in cultures of single hippocampal granule cells: a presynaptic form of plasticity. Neuron 16, 1147–1157.

Vaden, J.H., Banumurthy, G., Gusarevich, E.S., Overstreet-Wadiche, L., and Wadiche, J.I. (2019). The readily-releasable pool dynamically regulates multivesicular release. Elife 8.

Weisskopf, M.G., Castillo, P.E., Zalutsky, R.A., and Nicoll, R.A. (1994). Mediation of hippocampal mossy fiber long-term potentiation by cyclic AMP. Science 265, 1878–1882.

Xiang, Z., Greenwood, A.C., Kairiss, E.W., and Brown, T.H. (1994). Quantal mechanism of long-term potentiation in hippocampal mossy-fiber synapses. J Neurophysiol 71, 2552–2556.

Yang, Y., and Calakos, N. (2013). Presynaptic long-term plasticity. Front Synaptic Neurosci 5, 8.

Zalutsky, R.A., and Nicoll, R.A. (1990). Comparison of two forms of long-term potentiation in single hippocampal neurons. Science 248, 1619–1624.

Zucker, R.S., and Regehr, W.G. (2002). Short-term synaptic plasticity. Annu Rev Physiol 64, 355–405.

